# Scalable Agricultural Microbiome Sampling: Operational Definitions, Pooling Strategies, and Preservation Methods

**DOI:** 10.64898/2026.05.19.725853

**Authors:** Adam Ossowicki, Thom Griffioen, Enrichetta Mileti, Valentino Attanasi, Chris Hames, Victor J. Carrion, Ben Oyserman

**Affiliations:** Departamento de Microbiología, Facultad de Ciencias, Campus Universitario de Teatinos s/n, Universidad de Málaga, 29010 Málaga, Spain; Departamento de Protección de Cultivos, Instituto de Hortofruticultura Subtropical y Mediterránea ‘La Mayora’, Campus Universitario de Teatinos, Universidad de Málaga-Consejo Superior de Investigaciones Científicas (IHSM-UMA-CSIC), 29010 Málaga, Spain; Syngenta Crop Protection AG, Schaffhauserstrasse 101, Stein AG, Switzerland; Syngenta Biologicals, Via Cagliari 1 -Zona Industriale, 66041 Atessa (CH), Italy; Metagen Pty, Ltd., Gatton, Queensland, Australia; Institute of Biology, Leiden University, Sylviusweg 72, 2333 BE Leiden, The Netherlands; Department of Microbial Ecology, Netherlands Institute of Ecology (NIOO-KNAW), Wageningen, Netherlands; Syngenta Crop Protection, Rosentalstrasse 67, 4058 Basel, Switzerland

**Keywords:** Microbiome, Rhizosphere, Agriculture, Soil health, Amplicon sequencing, Sampling Standardization, Rhizocore, Sample preservation, Sample pooling

## Abstract

Scalable soil microbiome monitoring requires sampling methods that are reproducible across operators, field sites, and logistical constraints. Here, we evaluated three key methodological choices that commonly limit comparability in agricultural rhizosphere studies: how the rhizosphere sampling unit is operationally defined, sample pooling strategies, and preservation methods. We introduce the RhizoCore, a standardized root-zone soil core defined by core diameter, depth, position relative to the plant, and subsample volume, as a practical proxy for traditional rhizosphere sampling. The RhizoCore method captured more than 92% of the sequencing depth found in traditional rhizosphere samples, with differences limited predominantly to low-abundance taxa. Preservation methods significantly affected bacterial communities, while sample pooling showed greater impact on fungal diversity and substantially reduced within-group variability across all treatments. Despite these effects, differential abundance analysis revealed minimal compositional changes, with only a small fraction of microbial taxa significantly affected by either pooling or preservation method. Our findings demonstrate that the RhizoCore method provides a reproducible, and scalable approach for rhizosphere sampling that balances scientific rigor with practical field implementation, offering a framework for large-scale soil microbiome monitoring programs and for improving comparability among agricultural microbiome studies across diverse environmental conditions.

## Introduction

The soil microbiome is an important component of agroecosystem functioning, influencing nutrient cycling and uptake (Sokol et al., 2022), plant growth promotion (Glick, 2012), disease suppression (Gao et al., 2026), and soil structure (Hartmann and Six, 2022; Philippot et al., 2024). As the concept of soil health becomes generally accepted (Banerjee and van der Heijden, 2023; Kibblewhite et al., 2008; Lehmann et al., 2020), the challenge is shifting from demonstrating that microbiomes matter to measuring them reproducibly at scale (Doran, 2002). This is in part due to conflicting demands; the immense and growing pressures on the agricultural sector to increase production (Alexandratos and Bruinsma, 2012) balanced with the recognition that agricultural systems should develop technologies with minimal adverse environmental impacts (Pretty, 2008). As a result, it is increasingly imperative to implement hypothesis driven monitoring programs (Lindenmayer and Likens, 2010) for key parameters of Soil Health such as the microbiome.

One of the major hurdles to developing global soil microbiome monitoring programs is the lack of efficient, standardized, and reproducible sampling, sample preservation, and logistics that may be easily implemented globally by industry professionals, growers, citizen scientists, and researchers alike without undue economic or labor requirements. Such constraints are consequential for any monitoring program, and developing practical methodology that address them is crucial for their short- and long-term success. For example, simplified procedures for data collection outperform more quantitative yet resource consuming assessments of endangered species when resources are constrained (Joseph et al., 2006).Hence, establishing protocols that are both scientifically rigorous and practically feasible for field implementation may facilitate widespread adoption of standard soil microbiome monitoring. This, in turn, will support data-driven decision-making in agriculture, contribute to sustainable land management practices, enhance our understanding of soil ecosystem dynamics on a global scale, and develop new technologies that are compatible with, or may even enhance, the soil microbiome (Atwood et al., 2022).

In this study, we use scalable to mean a sampling and sample-handling workflow that can be applied consistently across large numbers of plants, fields, operators, and locations without a proportional increase in labor, specialist expertise, cold-chain requirements, or analytical cost. Scalability therefore encompasses operational throughput, logistical feasibility, and comparability of the resulting microbiome data.

The rhizosphere is defined biophysically, chemically and ecologically as the narrow region of soil directly influenced by roots physically, through secretions and microbial activity (Hinsinger et al., 2009) (Figure 1A). However, the rhizosphere’s boundaries are inherently dynamic through space and time (Kuzyakov and Razavi, 2019; Philippot et al., 2013; York et al., 2016) and difficult to delineate across time without specialized techniques. Most research concerning plant-associated microbiomes surveys the rhizosphere as the zone where the interactions and exchange between soil and plant microbiomes are concentrated. Often this highly active zone is compared to bulk soil which is not directly influenced by the plant roots and acts as a microbial bank with higher diversity but lower abundance and activity (Bulgarelli et al., 2013). Yet standardized protocols for rhizosphere extraction remain elusive. This lack of methodological consensus often limits the reproducibility and comparability of findings across independent investigations.

**Figure 1.**
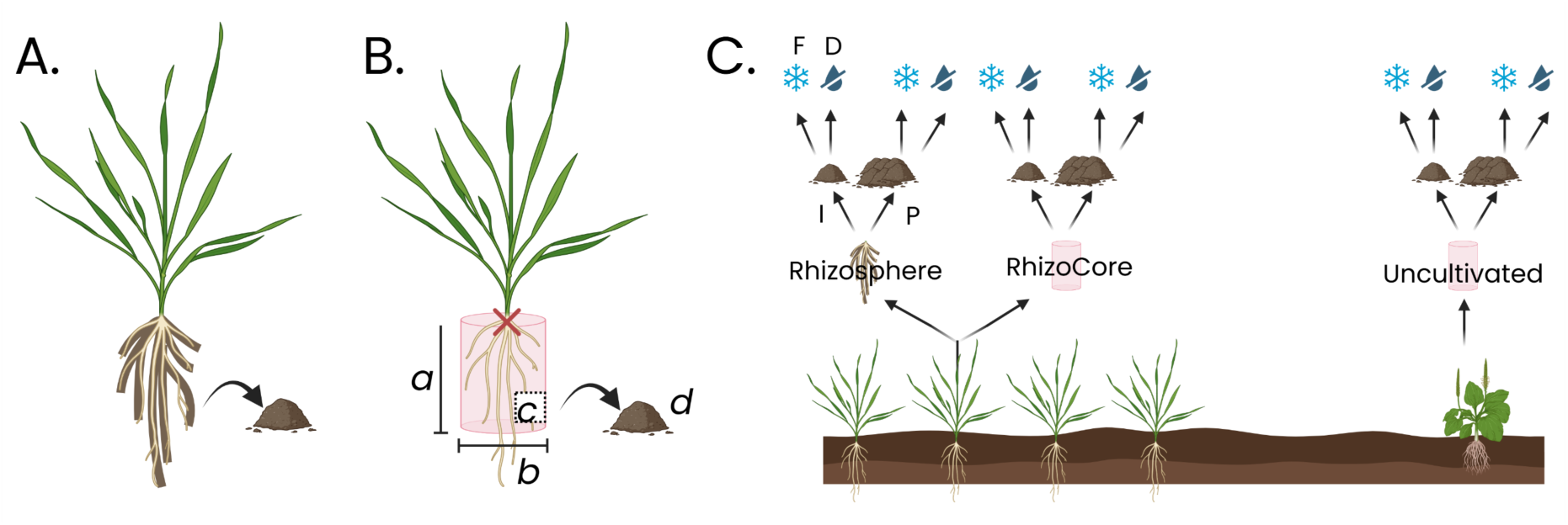
A. The rhizosphere as defined classically; the soil associated with the root zone that may be collected after shaking non-binding soil. B. The parameters defining the soil core: the diameter (b), the depth (a), the position of the soil core (c) and the volume of the sample (d). C. Experimental design in this study; three sampling zones: rhizosphere (R), RhizoCore and Uncultivated soil core as a bulk soil control, followed by pooling strategy: individual (I) and pooled *(P) and the type of preservation: freezing (F) and drying (D)*.

**Figure 2:**
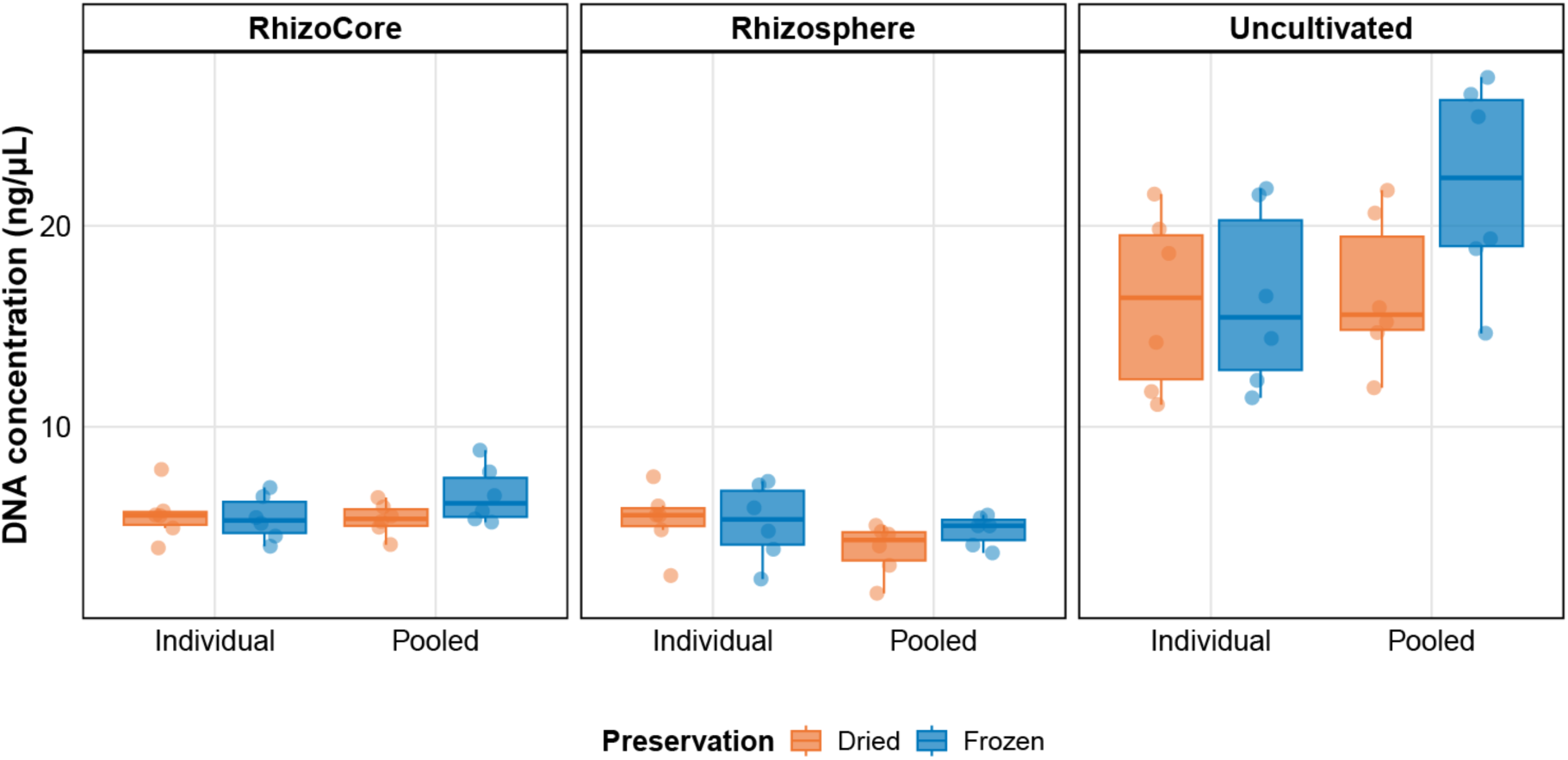
DNA concentrations obtained from soil samples across sampling zones, pooling strategies, and preservation methods.

The most common approaches require removing the plant from the ground, shaking off the excess of the soil (classified often as a bulk soil) and either brushing or scraping off the rhizosphere from the roots (Edwards et al., 2015; Ossowicki et al., 2020; Punt et al., 2026) or extracting it with a buffer (Bulgarelli et al., 2012; Chaparro et al., 2012; Lundberg et al., 2012) by shaking and optionally sonicating in it. Recently, Escudero-Martinez and colleagues systematically evaluated those “classical” methods of extracting rhizosphere for microbiome studies and metabolomics based on a greenhouse experiment (Escudero-Martinez et al., 2026). The rhizosphere extraction methods are generally not described in detail in the most or the present literature, leaving a lot of uncertainty in case of reproducing the study.

To address this, we introduce the operational definition of ‘rhizosphere core’ (RhizoCore) that maintains the ecological importance of the rhizosphere but improves the reproducibility and throughput of sampling. More specifically, we define the RhizoCore as a zone with a fixed diameter and depth, centered on and including the plant root system, which captures the rhizosphere and its immediate surroundings. This operational definition provides a universal sample unit by defining four parameters, the diameter of the soil core, the depth of the soil core, the position of the soil to sample and finally the volume of soil to sample (Figure 1B). These four parameters hence represent a minimal information for defining the exact sample location. By operationally defining the RhizoCore, we establish a trade-off of the precision of sampling the exact rhizosphere, with the convenience, speed, and reproducibility of the RhizoCore.

In addition, other major challenges are related to logistics and sampling preservation techniques which represent a hurdle to scale microbiome analysis. For example, pooled, or composite, soil microbiome samples have been used effectively for predictive purposes (Lutz et al., 2023; Romero et al., 2024). However, the impact of pooling samples on biodiversity estimates remains an important question to be addressed based on the aims of the study (Castle et al., 2019; Li et al., 2021). The decision to pool samples or analyze them individually may have implications for both the resolution of microbial community data and the practicality of large-scale soil microbiome assessments. Pooling may provide a cost-effective overview of field-level microbial diversity but may also obscure fine-scale heterogeneity, these hypotheses are not yet well tested in the literature. Conversely, individual sample analysis may offer detailed spatial information but at a higher processing cost for extraction and sequencing. Finally, another major topic of research is preservation techniques including freezing, drying, chemical preservation and the impact of temperature on unpreserved samples (Frøslev et al., 2023; Guerrieri et al., 2021; Smenderovac et al., 2024a). These studies have suggested that desiccating samples may be a promising alternative to freezing samples.

Here, we evaluate these three methodological issues together in an agricultural field setting. We compared traditional rhizosphere sampling, the RhizoCore approach, and an adjacent uncultivated reference core; tested individual samples against pooled composites; and compared flash freezing with silica drying. Using bacterial 16S rRNA and fungal ITS amplicon sequencing, we asked: (i) whether the RhizoCore captures the same dominant microbial signal as traditional rhizosphere sampling; (ii) how pooling and preservation affect diversity, community structure, and differential abundance; and (iii) whether these methodological effects are large enough to alter the ecological interpretation of sampling-zone differences. The broader goal is to provide a practical framework for reproducible and scalable agricultural microbiome sampling.

## Methods

### Sampling design and sample preservation

Soil samples were collected from a field in Stein Switzerland (47°33’11.0”N 7°58’16.5”E) growing Winter Wheat *Triticum aestivum* L. Variety AKTEUR (BBCH stage 30-31) on March 18th, 2024 Soil physical and chemical properties were analyzed by LUFA Speyer (Germany). The clay loam soil had a pH of 7.34, a total nitrogen concentration of 220 mg per 100 g total dry mass, an organic carbon content of 1.98% total dry mass, and a C/N ratio of 9; full soil-characterization data are provided in Table S1.

Three distinct zones were sampled as depicted in Figure 1 and were defined as follows:

Rhizosphere (Fig 1A) samples were obtained by gently shaking roots to remove loosely adhered soil. After this step, soil that was closely adhering to the plant roots was collected.
RhizoCore samples (Fig. 1B) were extracted using four parameters to operationally define the rhizosphere location: depth of 5 cm (*a*), core diameter of 10 cm (b), placement for sub sampling within the core parallel to the row crops (c), and finally volume of soil samples (10cm^3^).
Uncultivated – A Soil Core sample taken from an area adjacent to the field but not used for the cultivation with vegetation containing mostly grasses. The core was defined volume wise the same way it was defined in the RhizoCore samples.

From each individual sample, a subsample of approximately 10 cm^3^ was preserved immediately in two ways (below). The remaining material from each sample type was pooled, thoroughly mixed, and then subsampled six times to represent the pooled sampled.

Every individual and pooled sample was preserved in two different ways:

A. by immediate flash freezing 10 cm^3^ of sample in 15 ml conical tube, and
B. by adding 10 cm^3^ of sample to a small 3-inch by 5-inch Ziplock plastic bag with 10-gram silica gel desiccant (WiseDry).

As six replicates were collected for each combination of zone, preservation method, and pooling strategy, this resulted in a total of 72 samples that were taken for microbiome analysis (3 sampling zones x 2 preservation methods x 2 pooling strategies x 6 replicates, Fig. 1C).

This factorial sampling approach allows for comparison of microbial communities across different soil zones, preservation methods, and pooling strategies, providing a robust dataset for analysis of soil microbiome analysis.

### DNA Extraction G Sequencing

DNA extraction was performed on approximately 350 μL of soil by volume by Microsynth AG (Microsynth AG, Balgach Switzerland). Extraction, lysis, and DNA isolation was done according to manufacturer’s recommendations (ZymoBIOMICS DNA Mini Kit). Concentration of the isolated DNA was assessed with PicoGreen measurement (Quant-iT™ PicoGreen™ dsDNA Assay Kit, Thermo Fisher). From the extracted DNA, amplicons were generated in a two-step PCR approach. The first step amplified the 16S rRNA gene (V4 region), using the primers 515F (GTGYCAGCMGCCGCGGTAA) and 806R (GGACTACNVGGGTWTCTAAT) (Takahashi et al., 2014) (table S2). The second step amplified the ITS2 region using primers ITS3 (GCATCGATGAAGAACGCAGC) and ITS4 (TCCTCCGCTTATTGATATGC) (White et al., 1990) (table S3). Sequencing was performed by Microsynth AG, Balgach Switzerland using the Illumina NovaSeq 6000 platform for the 16S region, and the Illumina MiSeq for the ITS region. Both platforms ran 250bp paired end sequencing, with a target yield of 50 000 reads per sample. Sequences were demultiplexed and technical sequences were removed. Demultiplexing yielded 58 725 720 reads total.

### Bioinformatics

The sample sequencing files were processed to ASV’s using the nf-core/ampliseq pipeline version 2.16.1 (Straub et al., 2026). Methods of critical steps and follow up analysis are outlined below.

#### Amplicon Sequencing Processing

Data quality was evaluated with FastQC v0.12.1 (Andrews, 2023) and summarized with MultiQC v1.33 (Ewels et al., 2016). Cutadapt v5.2 (Martin, 2011) trimmed primers and all untrimmed sequences were discarded. Sequences that did not contain primer sequences were considered artifacts. Less than 18.6% of the sequences were discarded per sample and a mean of 97.6% of the sequences per sample passed the filtering.

#### DADA2 sequence processing and denoising

Adapter and primer-free sequences were processed sample-wise (independent) for both 16S and ITS amplicons using DADA2 v1.34.0 (Callahan et al., 2016). Filtered reads were trimmed by quality and length according to amplicon-specific parameters: maximum expected errors (maxEE = 2.2), truncation lengths (trunLen= 0), minimum read length (minLen = 50), and maximum ambiguous bases (maxN = 0). Reads not meeting these thresholds were removed. Samples with fewer than 12 500 reads were removed. The error models for forward and reverse reads were trained independently on the filtered data (nbases: 10^8^) using DADA2’s learnErrors(). Each sample was processed individually: dereplication of identical sequences (derepFastq()), denoising via the core DADA2 algorithm (dada()) using the error model, and paired-end merging (mergePairs()) with overhang trimming. Amplicon sequence variants (ASV’s) passing through all stages were retained in a sample-by-ASV abundance table. Chimeric sequences were identified and removed by consensus across samples (removeBimeraDenovo(method = “consensus”)).

#### Taxonomic assignment

Representative ASV sequences were assigned taxonomy with DADA2’s assignTaxonomy() function. For 16S dataset, the SILVA reference database (Chuvochina et al., 2026) version 138.1 was used. For ITS dataset, the UNITE reference database (Abarenkov et al., 2024) version 04.04.2024 was used. Bootstrap confidence thresholds were set to 60.

#### Data import and preprocessing

All analyses were performed in R v4.4.2 (R Development Core Team, 2019) using tidyverse v2.0.0 (Wickham et al., 2019) for data manipulation and phyloseq v1.52.0 (McMurdie and Holmes, 2013) for microbiome object handling. Taxa annotated as chloroplast or mitochondria, or lacking any annotation, were removed using subset_taxa(). Singletons were removed using prune_taxa() function.

#### Alpha diversity

Shannon index was calculated using QIIME2 and modelled using a linear model including all two-way interactions among Preservation method, Pooling strategy, and Sampling zone. Estimated marginal means and pairwise contrasts between Preservation methods were computed within each Pooling × Sampling_zone levels using the emmeans package (Lenth and Piaskowski, 2026), with Tukey’s HSD adjustment. Contrasts with p < 0.05 were considered significant. Results were visualized using ggplot2 R package.

#### Beta diversity and ordination

Beta diversity analysis was performed to assess the effects of sampling zone, preservation method, and pooling strategy on bacterial (16S) and fungal (ITS) community composition. Prior to analysis, raw amplicon count data were normalized using cumulative sum scaling (CSS) followed by log transformation, as implemented in the metagenomeSeq R package (Paulson et al., 2013). Bray-Curtis dissimilarity matrices were computed from normalized data using the phyloseq R package (McMurdie and Holmes, 2013). To test significant differences in community composition among groups, a permutational multivariate analysis of variance (PERMANOVA) was performed using the adonis2 function from the vegan R package (Oksanen et al., 2026). The model included the main effects of Sampling zone, Preservation, and Pooling, as well as all pairwise interactions.

To verify the assumption of homogeneous multivariate dispersion required by PERMANOVA, permutation tests of multivariate dispersions were conducted separately for each factor (Location, Preservation, and Pooling) using the betadisper and permutest functions in vegan. Community composition was visualized using Principal Coordinates Analysis (PCoA) based on Bray-Curtis dissimilarity, computed and plotted using phyloseq and ggplot2. Confidence ellipses (95%, t-distribution) were overlaid per group to aid visual interpretation.

For comparison of dispersion permutational analysis of multivariate dispersions (PERMDISP) was performed on Bray-Curtis dissimilarity matrices to test for homogeneity of variance among treatment groups. F-values and associated p-values were calculated using ANOVA on distances from samples to their group centroids. Main effects were tested for Sampling Zone (RhizoCore, Rhizosphere, Uncultivated), Pooling Status (Individual, Pooled), and Preservation Method (Frozen, Dried).

To test the effect of pooling strategy (pooled vs. individual samples) on community dispersion, we performed separate analyses for each sampling zone (RhizoCore, Rhizosphere, and Uncultivated). Statistical significance was assessed using permutation tests (999 permutations) implemented in the permutest function, which provides a non-parametric alternative to ANOVA that does not assume normality or homogeneity of variances.

#### Differential abundance

ANCOMBC version 2.12.0 was used to find differentially abundant taxa between treatments with significance threshold 0.05 and Holm p adjustment method (Lin and Peddada, 2020). Prior to the analysis the taxa showing zero or near-zero variance were removed using a custom script. The relative abundance (RA) was assessed by comparing total number of reads of specific taxa to the total number of reads in a dataset normalized by css log transformation, what was performed by using a custom R script.

## Results

### Sampling zone is the primary driver of DNA yield

An analysis of the DNA concentration revealed significant differences across sampling zones (F = 159.54, p < 0.001, Table S4). Uncultivated soil consistently yielded the highest DNA concentrations, ranging from approximately 15 to 22 ng/μL, markedly exceeding those of the agricultural RhizoCore and Rhizosphere zones, which both exhibited concentrations below 10 ng/μL. While the main effects of sample pooling (F = 1.75, p = 0.191) and preservation method (F = 3.42, p = 0.069) were not statistically significant, a notable interaction between zone and pooling method was observed (F = 3.22, p = 0.047). This interaction was particularly evident in Uncultivated soil samples, where frozen samples, especially when pooled, tended to yield higher DNA concentrations compared to dried samples. Interestingly, the Uncultivated soil samples also displayed greater variability in DNA concentration across treatments, potentially reflecting the heterogeneous nature of this environment. In contrast, RhizoCore and Rhizosphere zones showed minimal differences and consistent DNA yield across all the preservation methods and pooling strategies. These findings highlight that the effects of preservation and pooling methods may vary depending on the type of sample being studied. In summary, RhizoCore and rhizosphere samples did not differ significantly regardless of sampling methodology. Similarly, the difference between RhizoCore and Rhizosphere from Uncultivated were consistent. Regardless of sampling strategy, the Uncultivated samples always yielded the highest DNA concentrations, however statistical differences were observed with frozen pooled samples having highest DNA abundance.

### Alpha diversity confirms RhizoCore as a valid proxy for rhizosphere sampling

The analysis of microbial diversity defined by Shannon diversity index (Fig. 3, Table S5) reveals complex interactions between sampling zone, preservation method, and sample pooling on microbial community diversity. Across sampling zones: Rhizosphere, RhizoCore, and Uncultivated, no significant differences in the diversity index were observed for both bacteria and fungi. The preservation method shows a consistent, statistically significant effect (16S -p=0.0268, ITS – p=0.0113). Especially in Uncultivated sampling zone, dried samples typically exhibiting higher diversity than frozen samples in bacteria diversity but the opposite, lower in dried samples for fungi.

**Figure 3.**
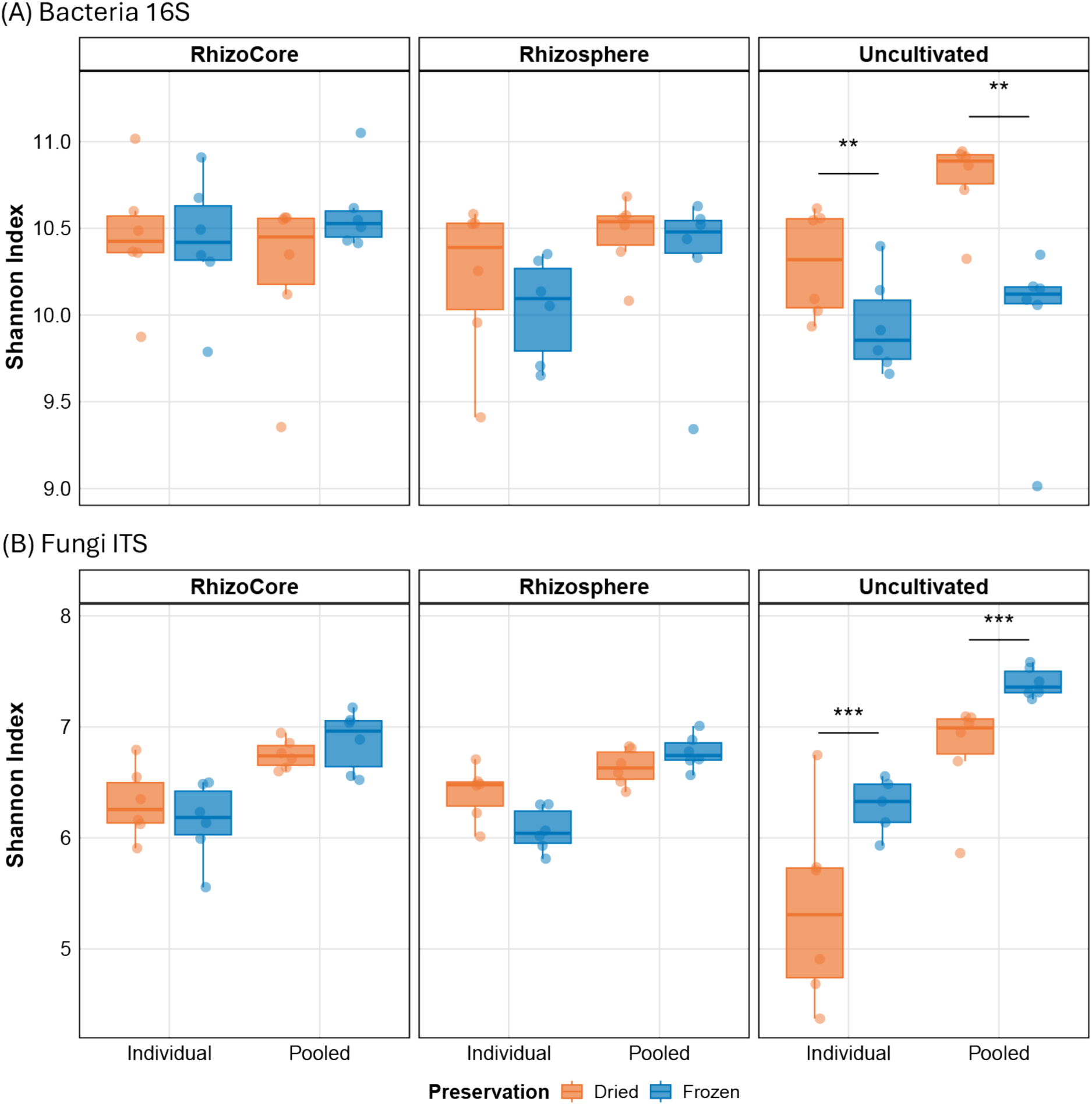
Bacterial (A) and fungal (B) diversity measured using Shannon index. Statistically significant differences marked with asterisk (linear model with Tukey-adjusted pairwise contrasts).

Sample pooling also influences diversity measures, with a significant effect in fungi (p < 0.0001) and close to reaching the significance threshold in bacteria (p = 0.0579). For both microbial groups the effect was the most pronounced in Uncultivated samples what probably reflects much higher heterogeneity of this environment. While in Uncultivated zone the highest diversity is recorded for fungi in frozen and pooled samples for bacteria it is in dried and pooled samples. This indicates that the pooling strategy generally is better to reflect the high diversity of uncultivated soil.

The considerable variability within groups underscores the heterogeneity of microbial communities even under similar treatment conditions. Nonetheless, pooling of the samples noticeably reduces the variability compared to the individual samples. These findings emphasize the critical importance of consistent sample handling and processing methods in soil microbiome studies to ensure accurate and comparable diversity assessments across different experimental conditions.

### Beta diversity confirms RhizoCore captures equivalent community composition to Rhizosphere sampling

For the bacterial community, the PERMANOVA analysis (Table S6A) revealed significant effects of not only sampling zone but preservation method and sampling zone on the microbiome. Sampling zone emerged as the strongest predictor, explaining 31.7% of the variation (R² = 0.317, F = 16.694, p < 0.001). Preservation method accounted for 2.7% of the total variation (R² = 0.027, F = 2.802, p = 0.011), indicating a measurable impact on community composition. While pooling showed smaller but also significant effect (R² = 0.019, F = 1.986, p = 0.045) it does not meet the requirement of data homogeneity evaluated by permutation test (p = 0.002).

In the fungal community, PERMANOVA (Table S6B) revealed the significant effect of pooling explaining 2.5% of the total variation (R² = 0.025, F = 2.78, p = 0.013) with the Sampling zone, though significant, and seeming to be the primary driver, (R² = 0.367, F = 20.786, p < 0.001) not passing the homogeneity criterion (p = 0.022). Notably, the interaction Zone:Pooling was statistically significant (p = 0.035).

For both bacteria and fungi, the PCoA plot (Fig. 4) and PERMANOVA results collectively provide a comprehensive view of the factors influencing microbial community composition in between tested treatments. Sampling zone emerged as the primary driver, nevertheless on the PCoA plot we can only clearly distinguish the Uncultivated zone while the Rhizosphere and RhizoCore are grouping together separating on axis capturing total variation of 31.7% and 35.8% for bacteria and fungi, respectively (Table S6 A and B). Based on preservation method, the samples do not form distinguishable clusters (Fig.4 A and B). Nevertheless, based on Pooling category (Fig.4 C and D) we can see two clear clusters of Pooled samples in both datasets, one common for Rhizosphere and RhizoCore and one for Uncultivated sampling zone what may be reflected in the statistically significant interaction Zone:Pooling (Fungal dataset). The data show that the pooling of samples can reduce the variation of the data.

**Figure 4.**
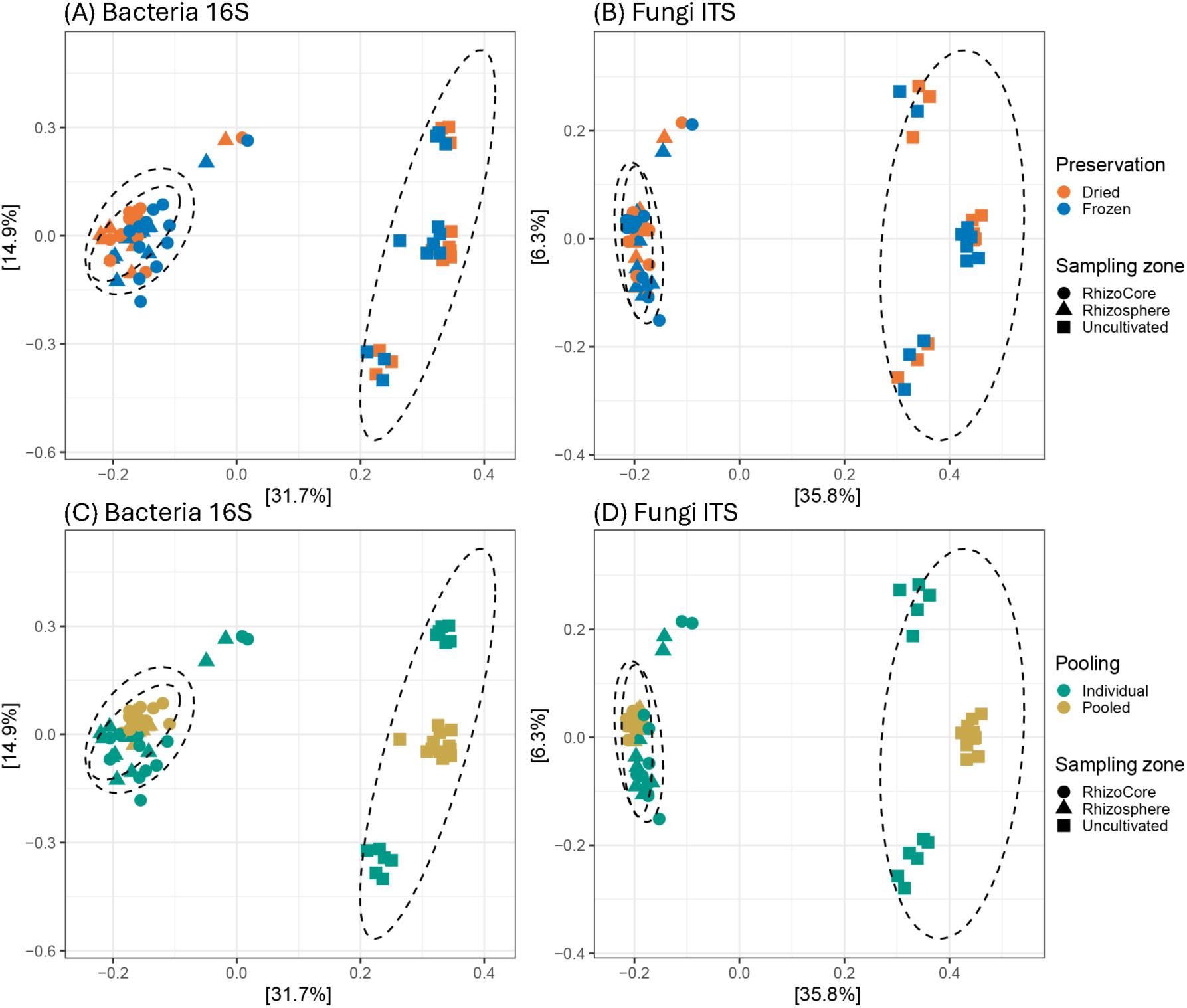
Principal Coordinates Analysis of bacterial (A & C) and fungal (B & D) microbial communities based on Bray-Curtis distances. The ellipses mark the strongest predictor – sampling zone. Plots A and B are colored by Preservation (red and blue) and plots C and D by Pooling (yellow and purple).

These findings highlight the complex interplay between sampling location, preservation method, and pooling in shaping soil microbial communities, emphasizing the importance of considering these factors in experimental design and data interpretation for soil microbiome studies.

### Pooling reduces within-group dispersion and improves community representation

Statistical assessment of within-group variability in soil microbial community composition revealed a significant impact of pooling on the dispersion of the bacterial community (F = 15.96, p = 0.0001) and nearly statistically significant in fungal community (F = 3.64, p = 0.060). Nevertheless, for both microbial groups pooling had a statistically significant impact of the dispersion in all three sampling zones (Table S7) what is visualized on Figure 5. This suggests that pooling has a significant impact on the observed community structure, potentially masking some of the natural variability present in soil microbial communities, this could be interpreted as presenting more consistent proxy of the microbiome for a given field or zone. The effect appears to be most pronounced in the Uncultivated zone, followed by the Rhizosphere, and then the RhizoCore zone.

**Figure 5.**
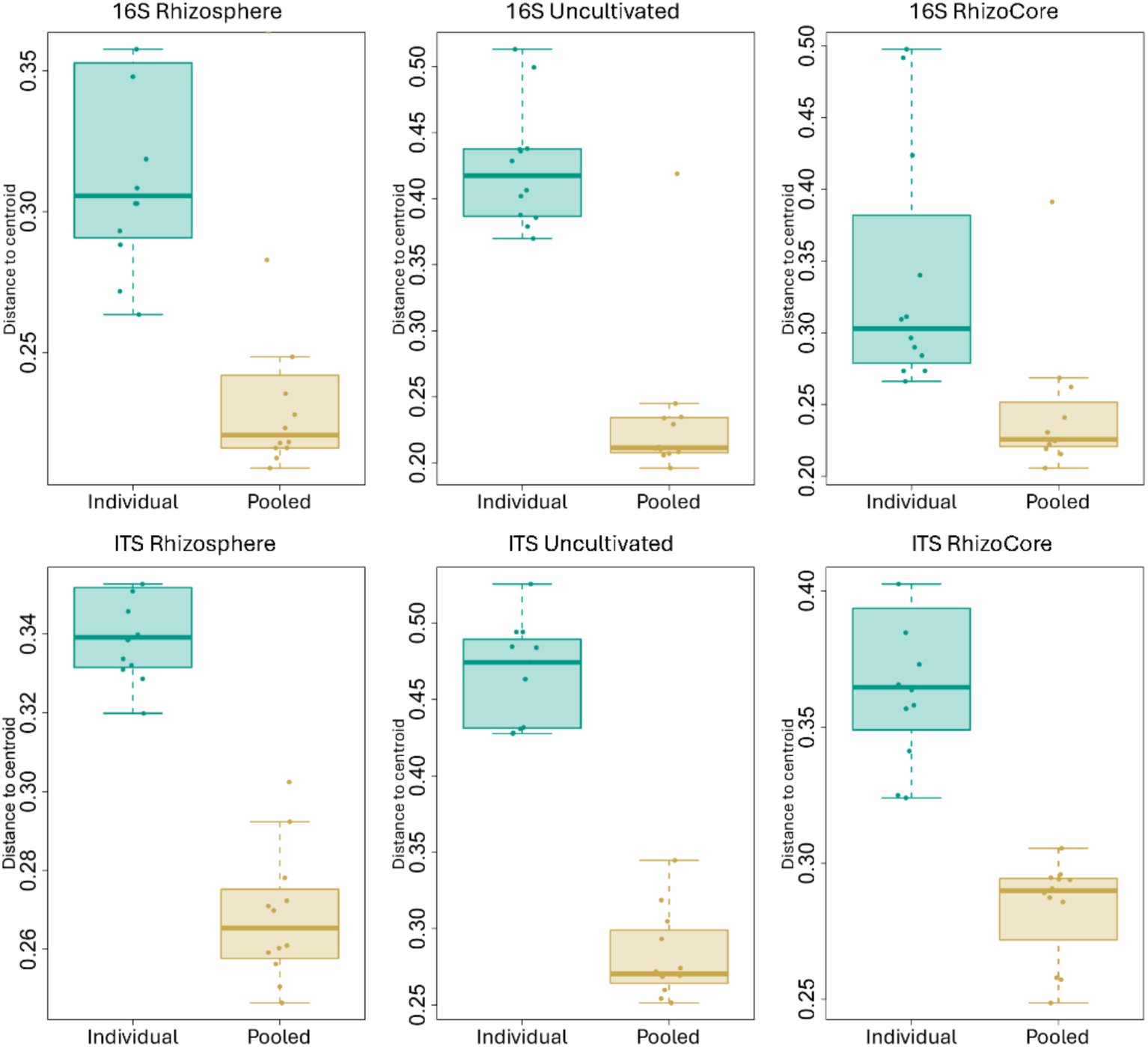
The effect of pooling samples across different sampling zones on the dispersion of microbiome as distance to centroid

### Compositional differences between RhizoCore and rhizosphere are limited to rare taxa

The differential abundance of taxa between treatments was tested using ANCOMBC2 on all taxonomy levels and finally at the ASV level. Only the statistically significant results (adjusted p-value < 0.05) with large biological effect (log fold change (lfc) >1) were considered relevant in this analysis. Often what makes biological interpretation challenging on different taxonomic levels is lack of annotation which directly relates to limited taxonomic resolution on sequence 16S or ITS fragments. Especially in the ITS dataset where between 41 and 48 % of ASVs were unannotated at the order, family and genus levels. The results are summarized in Table S8, further presented with more details in Table S9 and visualized in Figure 6.

**Figure 6.**
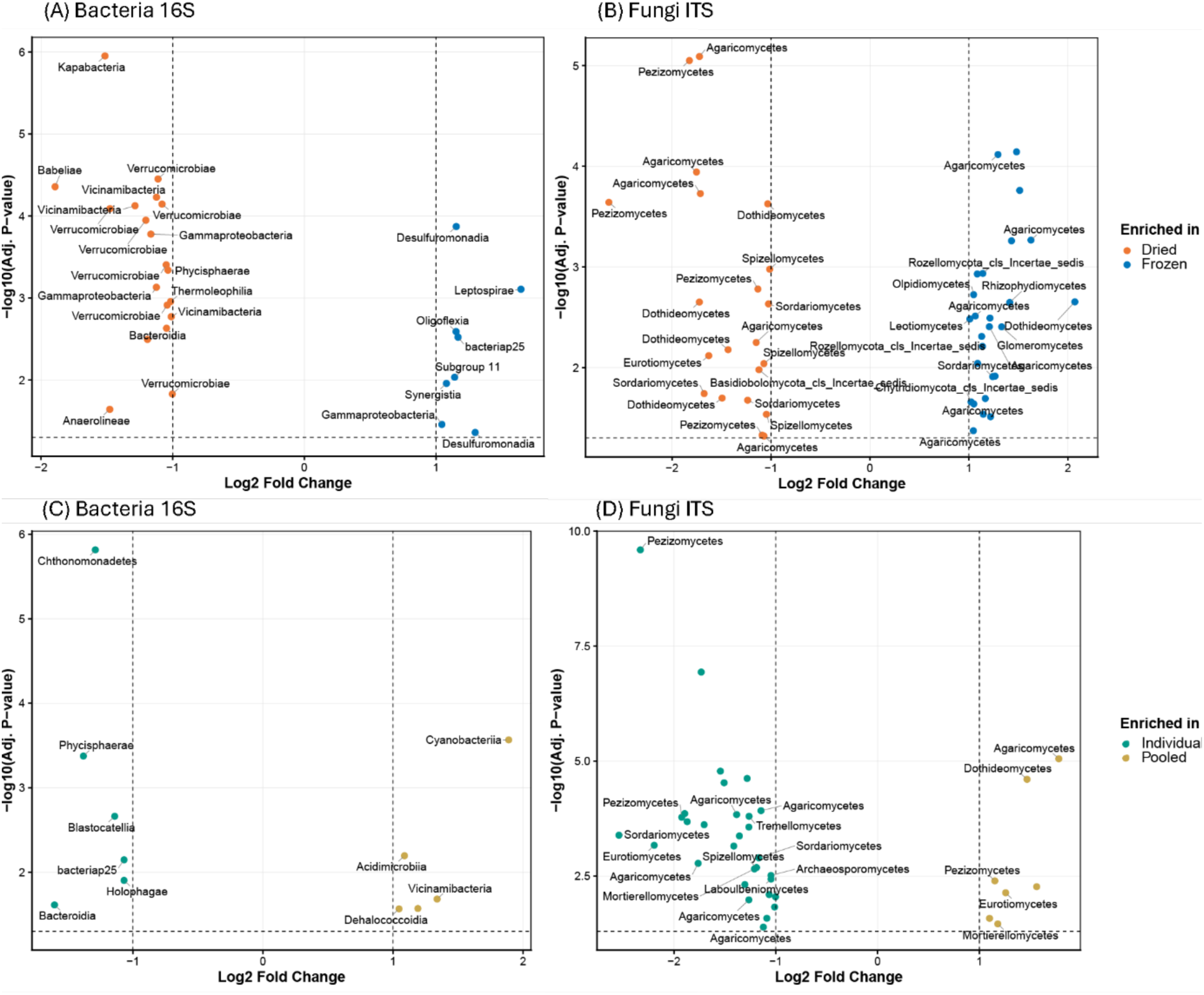
Volcano plots of statistically significant differential ASVs of bacterial (A & C) and fungal (B & D) microbial communities based on ANCOMBC2. Plots A & B show differences based on Preservation method and C & D based on Pooling. The ASVs are labeled with phylum for bacteria and class for fungi.

First the impact of pooling on the differential composition was evaluated. The results showed a minimal effect on both, bacteriome (16S) and mycobiome (ITS). Only 11 (< 0.09% RA – relative abundance) and 36 ASVs (< 2% RA) were differentially abundant between Individual and Pooled samples in 16S and ITS datasets, respectively.

In bacteria, pooling samples had an impact by enriching ASVs belonging to different phyla without clear taxonomic pattern, with different single ASVs of Acidobacteria enriched both in individual and in pooled samples. The highest taxonomic level where a difference was found was family level where one feature was enriched in pooled samples. The enriched feature belongs to Cyanobacteria family Phormidiaceae and was present at a low level (<0.04% RA). Also, the compound effect of pooling and sampling zone showed minimal impact with just one ASV enriched between Uncultivated treatments and the highest differential family Burkholderiaceae with also marginal share in the bacteriome (0.05% RA).

Fungal communities had more differentially abundant ASVs than bacteria, though the overall effect remained modest and did not extend to higher taxonomic levels. At the order level, only two taxa were enriched in Individual samples: Sebacinales and an unannotated order within class Dothideomycetes (combined <1% RA). Pooling samples had a disproportional effect with only 6 ASV’s enriched in Pooled samples and 30 in Individual what may suggest that some of the rare taxa diversity is lost due to pooling samples. The same as in bacterial community, in fungal community the compound effect of pooling and sampling zone is minimal with only two single ASVs enriched on each side of Uncultivated treatments, one from genus Marcelleina and the other from genus Cortinarius. This change account only for ∼0.08 % RA.

Such a low number of significant differences between individual and pooled Uncultivated samples is surprising because of the high diversity of those microbiomes but shows that pooling (regardless of sampling zone) has a marginal impact on the microbiome on compositional level in both bacteria and fungi.

In terms of the number of differentially abundant ASVs under the two DNA preservation procedures, freezing and drying in both bacteriome and mycobiome number of features was higher compared to Pooling. In the 16S dataset 27 differentially abundant ASVs (0.24 % RA) were identified and in ITS 47 ASVs (2.5 % RA).

The bacterial community showed 19 ASVs enriched in Dried treatment and only 8 in Frozen without clear taxonomic or functional pattern but dominated by gram negative cell wall structure except for one member of phylum Actinobacteria class Thermoleophilia enriched in Dried samples. On the higher taxonomic levels, a rare phylum Synergistota was enriched in Frozen samples, but it covered only 0.003% RA. The compound effect of Sampling zone and Preservation was marginal with only single ASVs or higher taxonomic groups with very low coverage enriched between treatments.

In the fungal community 47 ASVs were found differentially abundant in the Preservation category with 21 ASVs enriched in Dried samples and 26 in Frozen. The ASVs enriched in Dried samples are dominated by members of phylum Ascomycota. The difference in the highest taxonomic rank is a class in phylum Basidiobolomycota (0.02% RA) with unclear taxonomic placement. The interaction between Preservation and Sampling zone revealed even smaller changes with 3 ASVs differentially abundant between Rhizosphere samples.

Overall, the impact of preservation method on the differentially abundant taxa was bigger in fungi than in bacteria but still in both the groups it seems to alter mostly the very low abundant species. The performed differential analysis could not identify a major impact in any specific taxon under different treatment; in all the tested cases the effect was taxonomically variable.

The sampling zone Uncultivated was used in this study as a control highlighting the difference in microbiome between farming area where the microbiome is influenced by one crop and agricultural practices and the microbiome an area lacking agricultural pressure adjacent to the farm field. We can see the biggest differences between microbiomes in differentially abundant features between sampling zone Rhizosphere or RhizoCore and Uncultivated control what is expected based on the design of the study. In both the bacteriome and mycobiome large number of features on all taxonomic levels was found differentially abundant.

In bacterial community this accounts for 482 ASVs (13.1% RA) between RhizoCore and Uncultivated and 434 ASVs (13.7% RA) between Rhizosphere and Uncultivated. We can see striking changes in the bacteriome related to decrease in diversity already on the Phylum level with Elusimicrobiota and Entotheonellaeota enriched regardless of sampling zone and phyla Bdellovibrionota, Latescibacterota, and Verrucomicrobiota enriched in Uncultivated compared to Rhizosphere. Regardless of sampling technique, in both treatments Rhizosphere and RhizoCore when comparing with Uncultivated phyla Firmicutes, Nitrospirilla, Thermoplasmatota and WS2 are enriched. Also, the biggest log fold changes are observed between sampling zones six ASVs belonging to phyla Proteobacteria, Acidobacteriota and Bacteroidota decreased over 3 log fold between Rhizosphere or RhizoCore and Uncultivated. On the other hand, two Cyanobacteriia classes increased their relative abundance over 3 log fold in RhizoCore compared to Uncultivated.

Fungal community was also largely impacted between sampling zones, in Uncultivated samples we can identify 73 differentially abundant ASVs (13.7% RA) compared to RhizoCore and 77 ASVs (14% RA) comparing to Rhizosphere. The highest taxonomic group showing the changes was the class and in two comparisons, Rhizosphere against Uncultivated and RhizoCore against Uncultivated ten fungal classes were differentially abundant. Two of those Agaricomycetes and Archaeosporomycetes were both enriched in Uncultivated treatments while the first one has a significant presence in the ITS dataset accounting for almost 8.6% of the total RA and decreasing the abundance in both RhizoCore and Rhizosphere more than 3.2 log fold. Also three fungal classes, Taphrinomycetes, Zoopagomycetes and, Dothideomycetes were significantly enriched in both RhizoCore and Rhizosphere compared to Uncultivated. The biggest log fold changes of single features are identified between Uncultivated treatment and Rhizosphere or RhizoCore are pointing into wild mushrooms therefore suggesting that they are related to land use, the species *Inocybe griseovelata* and different members of the family Thelephoraceae decreased their abundance by 6-7 log fold.

While the differences in composition between Uncultivated treatments and other sampling zones reflect the differences between agricultural and non-agricultural setup comparing treatments Rhizosphere and RhizoCore shows the impact that using different sampling techniques has on the final microbiome.

The number of differentially abundant taxa or ASVs was very small, the analysis found only three differential ASVs (0.02% RA) in the 16S dataset and seven ASVs (0.6% RA) in ITS. In the bacteriome the only significant features (ASVs and order) occur within the phylum Planctomycetota, nevertheless different features were enriched on both sides, only one ASV is enriched exclusively in Rhizosphere and belongs to genus Hydrogenophaga.

The mycobiome shows similar minor changes almost exclusively within phylum Ascomycota with differential features within classes Sordariomycetes, Eurotiomycetes and Dothideomycetes. Only higher taxonomy level showing significant changes was the order Sebacinales enriched in RhizoCore (> 0.08% RA).

To further investigate the impact of sampling zone additional comparison was undertaken. The two microbiomes resulting from classical rhizosphere sampling (R) and one using the RhizoCore approach were very similar to each other what as has been already shown in the alpha diversity analysis (Fig.3), beta diversity analysis (Fig. 4) and differential abundance analysis. To highlight the similarities the shared and unique ASVs passing a 50% prevalence filter were plotted on a Venn diagram together with the share in the total sequencing depth between treatments.

Comparison revealed that in the bacteriome both locations share 74 % individual ASVs but those shared ASVs account for the 92.2% of the sequencing depth. The exclusive part of the microbiome consists of 6% for the Rhizosphere samples and 20% for RhizoCore in terms of individual ASVs that is 1.62 % and 6.13 % of the total sequencing depth respectively. The mycobiome shows very similar pattern with around 69% shared ASVs accounting for 94,75% of total RA leaving the remaining 2.44% exclusive in RhizoCore and 2.85% in the Rhizosphere.

The number of taxa detected exclusively in sampling zones regardless of the amplicon sequenced (16S or ITS) with relatively low share in the total sequencing volume in those samples most probably is a result of larger fraction of soil more distant from the root with relatively smaller impact of the root chemistry in the sample RhizoCore, still the exclusive share consists mostly of low abundant to rare taxa. This hypothesis is supported by the differential abundance analysis resulted only in 3 consistently differential ASVs in 16S dataset and only 7 in ITS between Rhizosphere and RhizoCore (Tables S8 and S9).

## Discussion

The rhizosphere is one of the basic concepts of plant–microbiome interaction studies and has been defined for more than 100 years as “the soil compartment influenced by plant roots” (Bakker et al., 2013). However, how much soil is “influenced” is ultimately a subjective determination made by the sampler during harvest. This poses a major hurdle, as sampling units based on subjective definitions are difficult to replicate (Mörsdorf et al., 2015) and are particularly problematic in microbial ecology, where community composition can change sharply over millimeter-to-micron scales. Despite this challenge, the fundamental task of defining a standardized sampling unit for rhizosphere studies has not been thoroughly explored. We therefore propose establishing a formal *operational definition* of the rhizosphere sampling unit as an important step toward advancing plant–microbiome research.

We introduce the RhizoCore as this operational definition, specified by the distance from the stem (a), depth (b), the orientation/position of the soil core, when necessary (c), and the volume of soil sampled (d) (Figure 1b). Importantly, the RhizoCore is intended as a reproducible sampling unit that supports scale-up across operators, sites, and sampling campaigns through standardized geometry and rapid field collection. The RhizoCore is reproducible because it provides explicit geometric criteria that can be consistently applied across operators, sites, and time points. At the same time, because the ecological rhizosphere is defined by root influence rather than geometry, RhizoCore should be understood as an operational proxy that produces a controlled, volume-averaged estimate of the root-zone microbiome. Thus, RhizoCore is not intended to replace the rhizosphere as an ecological concept, but to provide a standardized and scalable method to measure it in practice.

Beyond introducing the RhizoCore as a reproducible and formal definition of the sampling unit, this study evaluates two additional methodological factors often used in microbiome research, the effects of sample pooling strategies (Lutz et al., 2023; Romero et al., 2024), and the impact of various preservation methods (Frøslev et al., 2023; Guerrieri et al., 2021; Smenderovac et al., 2024b). By addressing these issues together, we aim to develop a reproducible and scalable soil-microbiome workflow across diverse agricultural settings. This approach is required to bridge the gap between scientific understanding and practical application in the field, allowing for more effective integration of microbiome data into agricultural decision-making processes (Doran, 2002; Pretty, 2008). Our research not only contributes to the standardization of soil health assessments but paves the way for large-scale, comparative studies that can inform sustainable land management practices on a global scale (Lindenmayer and Likens, 2010).

The need for standardized sampling units is not unique to rhizosphere research. Estimation of diversity is a central consideration across ecological disciplines, and methods must balance accuracy, efficiency, and reproducibility. A classic example is the use of quadrats in terrestrial ecology, which standardized sampling of plant communities by defining fixed areas for observation and measurement (Barbour et al., 1999). This approach significantly enhanced the comparability of vegetation studies across different ecosystems and research teams, and has also been adopted for aquatic systems (Vaughn, 1997).

Our results demonstrate that the RhizoCore method provides diversity estimates comparable to those obtained from traditional rhizosphere sampling techniques. Importantly, the RhizoCore method exhibited consistent performance across different preservation and pooling treatments, with minimal variations in DNA yield between RhizoCore and Rhizosphere zones. Nevertheless, because RhizoCore is a standardized geometric proxy, it may not capture the full complexity of the rhizosphere. Despite that, most of the sequenced microbiome overlapped between RhizoCore and Rhizosphere samples (92.23% for bacterial 16S and 94.75% for fungal ITS; Figure 7), while differences were largely confined to low-abundance taxa. Furthermore, as demonstrated in the differential abundance analysis, RhizoCore and Rhizosphere samples had minimal statistically significant differences. For bacteria, only 3 ASV were differentially, and none at family level and above (Table S7). For fungi, only 7 ASVs were differentially abundant, and none at class level and above (Table S7). Equally important, comparisons to an outgroup yielded similar conclusions: RhizoCore vs. Uncultivated showed 482 differentially abundant ASVs, while Rhizosphere vs. Uncultivated showed 434, indicating that choice between RhizoCore and traditional rhizosphere sampling method is unlikely to alter broad ecological interpretations when contrasting to a distinct reference condition. This aligns with Escudero-Martinez et al. (2026), who reported that despite protocol-specific differences in rhizosphere microbiota composition, approximately 75% of taxa distinguishing the rhizosphere from unplanted soil, analogous to our Uncultivated control, were consistently recovered across sampling approaches, although that study was conducted in greenhouse pots, our findings extend this observation to a field setting with additional evaluation of preservation and pooling strategies

**Figure 7.**
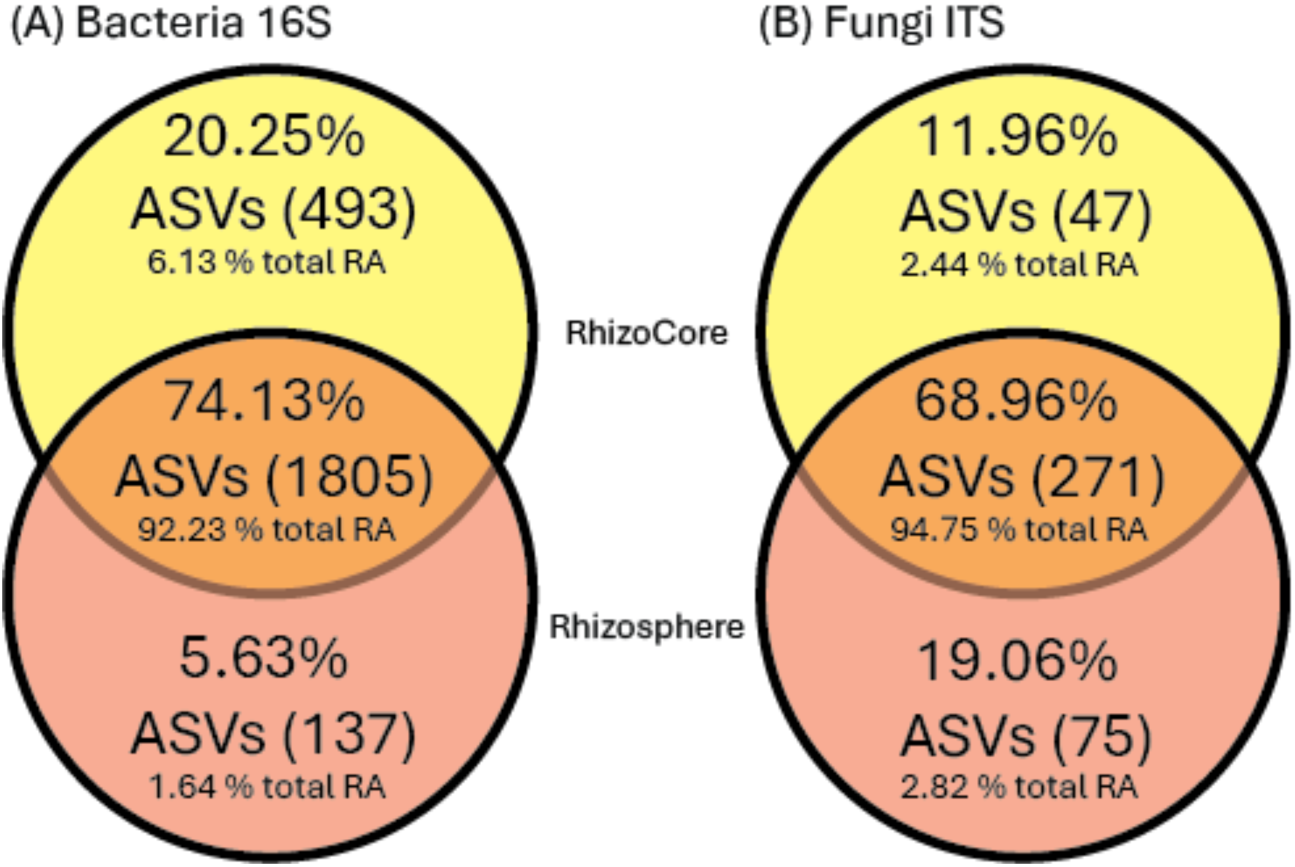
Venn diagram showing the difference in the number of ASVs passing a 50% prevalence filter between Rhizosphere and RhizoCore treatments.

The RhizoCore framework is both versatile and rapid. This allows for adaptation across diverse plant architectures and experimental settings (Fig. 8). While annual crops may be sampled with the core centered over the stem as done in this study (Fig. 8B), the method is easily adjusted for perennials to avoid destructive sampling (Fig. 8C). In such cases, the core can be placed at a fixed lateral distance from the stem, with the sample position within the core adjusted appropriately. Furthermore, the RhizoCore offers efficiency for greenhouse studies (Fig. 8D). Traditional rhizosphere sampling requires meticulous manual separation or root washing, can take 10 minutes or more per plant. For large studies, such as those with many genotypes, this requires multiple days of harvest (Oyserman et al., 2022). In contrast, the RhizoCore procedure is completed in approximately one minute, allowing experiments involving hundreds of units to be harvested in a single day rather than across a multi-day window. By consolidating the harvest time, researchers may eliminate a confounding variable of experimental noise (e.g. ‘Harvest Day’).

**Figure 8:**
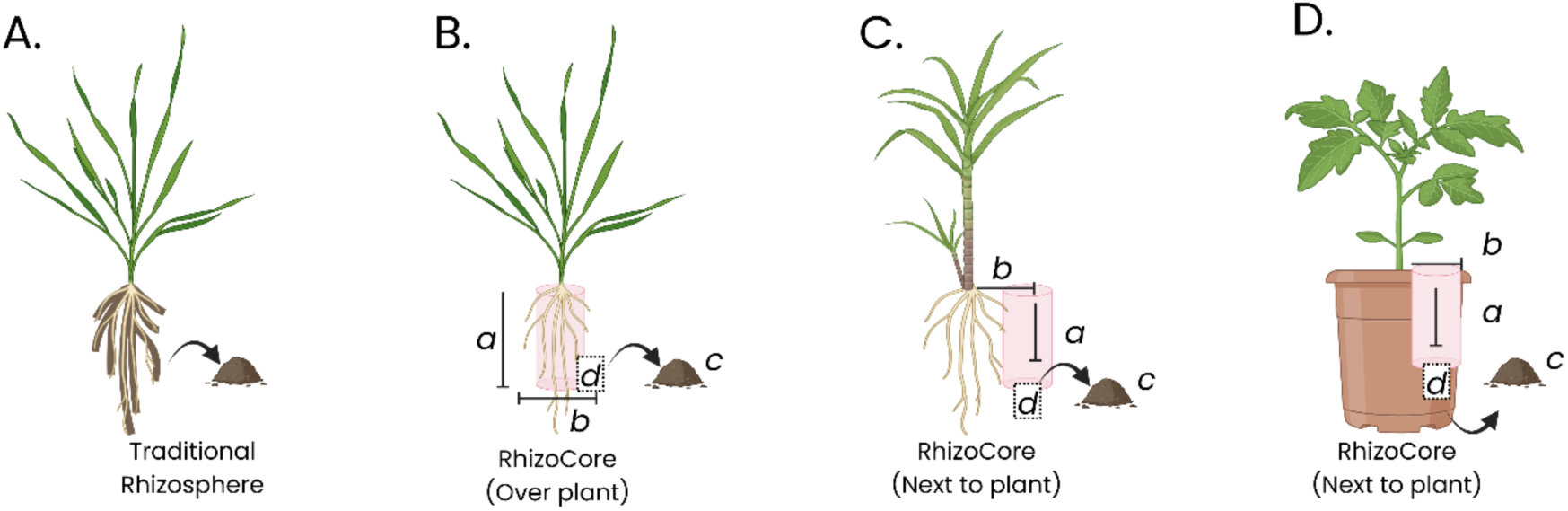
(A) Classic definition of the rhizosphere (B) Application of the RhizoCore in annual crops, where the core is centered over the stem. Requiring destructive sampling (C) Adapted application for perennial crops or trees to facilitate non-destructive sampling; the core is placed at a fixed lateral distance from the stem, with the internal sample position adjusted for spatial consistency. (D) Implementation in high-throughput greenhouse studies. By reducing sampling time from approximately 10 minutes (traditional manual separation/root washing) to 1 minute per plant, the RhizoCore allows for the harvest of hundreds of units in a single day which may be beneficial even in non-field conditions.

We also investigated the effects of sample pooling and preservation on diversity estimates. The impact of pooling and preservation varied between bacterial and fungal communities. For bacteria, sampling zone was the primary driver of diversity (explaining 31.7% of variation), while preservation method also significantly impacted diversity estimates (2.7% of variation) (Table S5A). Pooling showed a small but statistical significance, however, did not pass the permutation data homogeneity test making the difference statistically difficult to interpret. For fungi, both sampling zone (36.7% of variation) and pooling (2.5% of variation) significantly affected diversity estimates, with a notable interaction between pooling and Sampling zone (4.46% of variation) (Table S5B). Pooling generally increased microbial diversity, especially in frozen samples, and reduced variability as evidenced by narrower confidence intervals. Moreover, effect of pooling in all the sampling zones was significant in considering the dispersion of the data reducing the variability within and between the replicates. The reduction in variability may come at the cost of obscuring natural heterogeneity in soil microbial communities. Pooling effects impacted only rare taxa (<0.09% RA bacteria, <2% RA fungi out of total sequencing depth). These findings demonstrate that while both preservation and pooling introduce some bias, their effects are largely confined to rare taxa, suggesting either method is suitable for capturing core community composition. This interpretation is consistent with a recent study of arbuscular mycorrhizal fungi, which similarly found that pooling individual samples leads to missing rare arbuscular mycorrhiza taxa and using drying rather than freezing did not substantially alter the main biological conclusions, despite some treatment-dependent variation in community estimates (Frew et al., 2026).

Literature suggests that both dominant (Jiao et al., 2019) and rare taxa (Xiong et al., 2021) contribute to overall microbial community function and stability. Most of the studies comparing the impact of storage and preservation methods on microbiomes focus on gut and fecal samples. Nevertheless, the available literature investigating soil samples shows that the procedure of the storage and especially the time before sequencing has an impact on the diversity and composition of the final microbiomes (Guerrieri et al., 2021; Lauber et al., 2010; Rubin et al., 2013). However, studies focusing on rare taxa should carefully consider these methodological impacts.

Future research should focus on further optimizing and standardizing soil sample handling protocols to enhance reproducibility and sample quality. For example, one area for future investigation is determining precise desiccant-to-sample ratios for efficient drying, considering that silica gel can absorb approximately 30% of its mass in water. Additionally, streamlined logistics should be further investigated to minimize processing time between sampling and DNA extraction. One approach that may be considered would be to implement a hybrid preservation method where samples are initially dried during transit and then frozen upon arrival at processing facilities. In practice, we have observed that transit times globally may be between 1 day and 1 week using priority mail. This could reduce the period samples are left ‘unfrozen’, which we continue to assume as the gold standard in preservation, while waiting for extraction.

Finally, although this study focused on rhizosphere sampling, the same standardization principles could be extended to other plant-associated microbiomes, including phyllosphere and endosphere communities. However, these environments present unique challenges, such as lower biomass in the phyllosphere and the need for surface sterilization in endosphere sampling, which require dedicated methodological investigations. Understanding how these different preservation approaches affect low-abundance taxa and functional groups across various plant tissues will be crucial for developing comprehensive, standardized protocols for field based plant-microbiome research.

## Conclusions and Recommendations

Reproducibility is a central requirement of the scientific process and has come under increased scrutiny in recent years. A key contributor is the lack of standardized experimental methods and incomplete reporting of critical details which prevents independent researchers from accurately replicating published protocols (Baker, 2016; Bindels et al., 2025). This methodological variability has led to substantial waste of research resources and delays in translating basic discoveries into applications. This is especially important in areas that deal with complex multi-variable interactions such as agricultural microbiomes which vary considerably through space and time.

This study addresses three fundamental challenges in soil microbiome research by evaluating: (1) an operational definition of rhizosphere sampling through the RhizoCore approach, (2) the impact of sample pooling, and (3) the effects of preservation methods. Our findings demonstrate that the RhizoCore method provides a reliable alternative to traditional rhizosphere sampling, yielding comparable microbial diversity estimates while improving the reproducibility of the sampling unit.

RhizoCore captured 92.2% of the bacterial sequencing depth found in traditional rhizosphere samples, with differences primarily occurring in low-abundance taxa. This high overlap suggests that the RhizoCore method effectively captures the core rhizosphere microbiome while providing a standardized sampling protocol that can be consistently implemented across different studies and field conditions. Importantly, the RhizoCore is intended as an operational proxy for the ecologically defined rhizosphere, not a replacement of the concept.

Pooling and preservation had distinct, community-specific effects. Sampling zone was the primary driver of community composition for both bacteria (31.7% of variation) and fungi (36.7% of variation), while preservation significantly affected bacterial communities (2.7%) and pooling had a stronger effect on fungal communities (2.5%). Pooling reduces variability across treatments, but may also mask natural heterogeneity, particularly in uncultivated soils. Nevertheless, because sampling zone remained the dominant determinant of community structure, these methodological choices were generally secondary to the ecological signal of interest.

The effects of preservation methods were largely minor, however showed taxon-specific effects, with certain bacterial phyla (e.g., Firmicutes, Desulfobacterota) and fungal groups responding differently to preservation techniques. These findings emphasize the importance of consistent preservation methods when comparing microbial communities across studies and understanding potential biases that are introduced by sample handling.

Overall, our results provide a framework for standardized soil microbiome sampling that balances scientific rigor with practical field implementation. The RhizoCore method, combined with appropriate pooling and preservation strategies, enables reproducible and scalable sampling suitable for large-scale soil microbiome monitoring and supports the integration of microbiome data into soil health assessment and sustainable agricultural management.

## Data availability

The raw sequencing data (16S rRNA and ITS amplicon sequences) generated in this study have been deposited in the European Nucleotide Archive (ENA) at EMBL-EBI under accession number PRJEB113196. Data will be publicly available upon publication.

## Acknowledgements

We would like to thank Léo Britschu, Philippe Butz, Alice Giletta, Morgane Klink and Jan Werthmueller for their support.

## CRediT authorship contribution statement

**A.O.** -Writing – review & editing , Writing – original draft, Visualization, Validation, Formal analysis, Data curation; **T.G.** -Writing – review & editing, Formal analysis, Data curation; **E.M.** -Writing – review & editing, Formal analysis, Visualization; **V.A.** -Writing – review & editing, Formal analysis; **C.H.** -Writing – review & editing, Formal analysis; **V.J.C.** -Writing – review & editing, Funding acquisition; **B.O.** -Writing – review & editing , Writing – original draft, Methodology, Investigation, Funding acquisition, Conceptualization, Project administration

