## Supplementary material1 for "Scalable Agricultural Microbiome Sampling: Operational Definitions, Pooling Strategies, and Preservation Methods"

Ossowicki et al. 2026

### Table S1. Physical and chemical characteristics of the soil used in the study

| **22-Dec** | **uni. (LUFA Speyer)** | **Stein filed** |
| --- | --- | --- |
| **Dry Mass** | **%FM** | 89.4 |
| **Soil Type** |  | Clay loam |
| **pH** |  | 7.34 |
| **C** | **% DM** | 1.98 |
| **Total N** | **mg/100g DM** | 220 |
| **C/N** |  | **9** |
| **P2O5** | **mg/100g DM** | 11 |
| **K2O** | **mg/100g DM** | 9 |
| **Mg** | **mg/100g DM** | 16 |
| **B** | **mg/1kg DM** | 0.46 |
| **S** | **mg/l** | 2.03 |
| **S** | **kg/ha (30cm)** | 39.2 |
| **Cu** | **mg/1kg DM** | 3.62 |
| **Mn** | **mg/1kg DM** | 25.4 |
| **Zn** | **mg/1kg DM** | 5.73 |
| **KCl** | **mg/100g DM** | 79 |
| **Fe** | **mg/1kg DM** | 66.7 |
| **Clay (%)** | **<0.002mm** | 29.4 |
| **Slit (%)** | **0.002-0.050mm** | 45.6 |
| **Sand (%)** | **0.05-2mm** | 25 |
| **CEC** | **cmol/kg DM** | 20.8 |

### *DM – dry matter, FM – fresh matter

### Table S2. Sequencing information 16S amplicon sequencing

| 16S  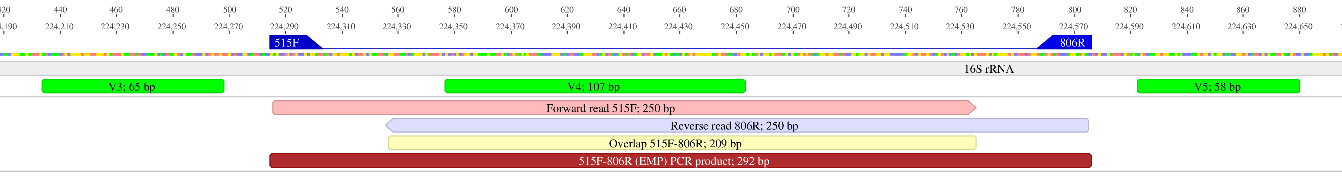 | |
| --- | --- |
| Sequencing provider | Microsynth AG Balgach |
| Targeted region | 16S V4 (~300 bp) |
| Target read pairs per sample | 50 000 |
| Forward primer | 515F  GTGYCAGCMGCCGCGGTAA |
| Reverse primer | 806R  GGACTACNVGGGTWTCTAAT |
| Sequencing method | Illumina NovaSeq 6000  Illumina DNA prep, SP flow cell |
| Fragment size | 2 x 250bp |

### Table S3. Sequencing information ITS amplicon sequencing

| ITS  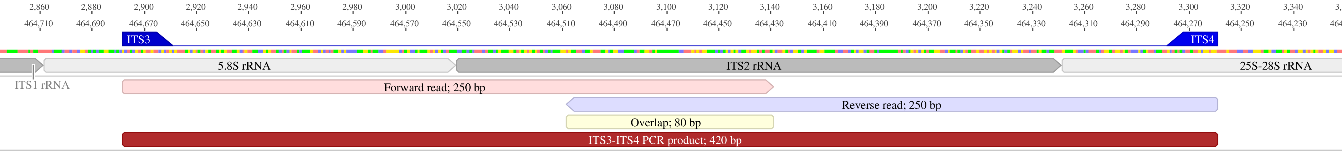 | |
| --- | --- |
| Sequencing provider | Microsynth AG Balgach |
| Targeted region | ITS2 (300~400 bp) |
| Target read pairs per sample | 50 000 |
| Forward primer | ITS3  GCATCGATGAAGAACGCAGC |
| Reverse primer | ITS4  TCCTCCGCTTATTGATATGC |
| Sequencing method | Illumina MiSeq |
| Fragment size | 2 x 250bp |

### Table S4. Summarized statistics DNA extraction concentrations. ANOVA table.

| Source | df | Sum Sq | Mean Sq | F | p-value |
| --- | --- | --- | --- | --- | --- |
| SamplING ZONE | 2 | 2510,98 | 1255,49 | 159,54 | < 0.001 *** |
| Preservation Method | 1 | 26,91 | 26,91 | 3,42 | 0,069 |
| Pooling Strategy | 1 | 13,77 | 13,77 | 1,75 | 0,191 |
| ZONE × Preservation | 2 | 21,03 | 10,51 | 1,34 | 0,271 |
| ZONE × Pooling | 2 | 50,63 | 25,31 | 3,22 | 0.047 * |
| Preservation × Pooling | 1 | 28,97 | 28,97 | 3,68 | 0,06 |
| Three-way Interaction | 2 | 15,85 | 7,92 | 1,01 | 0,371 |
| Residuals | 60 | 472,16 | 7,87 |  |  |

### Table S5. Summarized statistics bacteria (a) and fungi (b) alpha diversity as Shannon index

| **(A)Bacteria 16S** | | | | | | |
| --- | --- | --- | --- | --- | --- | --- |
| **ANOVA** | | | | | | |
| Term | Df | Sum Sq | Mean Sq | F value | Pr(>F) | Significance |
| Preservation | 1 | 0,7058 | 0,7058 | 5,14 | 0,0268 | * |
| Pooling | 1 | 0,5124 | 0,5124 | 3,73 | 0,0579 | ns |
| Sampling_zone | 2 | 0,5098 | 0,2549 | 1,86 | 0,1646 | ns |
| Preservation:Pooling | 1 | 0,0023 | 0,0023 | 0,02 | 0,8980 | ns |
| Preservation:Sampling_zone | 2 | 1,6508 | 0,8254 | 6,02 | 0,0041 | ** |
| Pooling:Sampling_zone | 2 | 0,2966 | 0,1483 | 1,08 | 0,3456 | ns |
| Residuals | 62 | 8,5072 | 0,1372 | NA | NA |  |

| emm contrast results Contrast \| Pooling \| Location \| Estimate \| SE \| df \| t.ratio \| p.value; Dried - Frozen \| Individual \| RhizoCore \| -0,1683 \| 0,1746 \| 62 \| -0,9638 \| 0,3389; Dried - Frozen \| Pooled \| Rhisosphere \| -0,1458 \| 0,1746 \| 62 \| -0,8352 \| 0,4068; Dried - Frozen \| Individual \| RhizoCore \| 0,1570 \| 0,1746 \| 62 \| 0,8989 \| 0,3722; Dried - Frozen \| Pooled \| Rhisosphere \| 0,1794 \| 0,1746 \| 62 \| 1,0276 \| 0,3081; Dried - Frozen \| Individual \| RhizoCore \| 0,5717 \| 0,1746 \| 62 \| 3,2740 \| 0,0017; Dried - Frozen \| Pooled \| Rhisosphere \| 0,5942 \| 0,1746 \| 62 \| 3,4026 \| 0,0012 | | | | | | |
| --- | --- | --- | --- | --- | --- | --- |
| **(B) Fungi ITS** | | | | | | |
| **ANOVA** | | | | | | |
| Term | Df | Sum Sq | Mean Sq | F value | Pr(>F) | Significance |
| Preservation | 1 | 0,9681 | 0,9681 | 7,1402 | 0,0097 | * |
| Pooling | 1 | 10,5338 | 10,5338 | 77,692 | 0 | *** |
| Sampling_zone | 2 | 0,0744 | 0,0373 | 0,2748 | 0,7606 | ns |
| Preservation:Pooling | 1 | 0,0915 | 0,0915 | 0,6752 | 0,4144 | ns |
| Preservation:Sampling_zone | 2 | 2,6821 | 1,340 | 9,8909 | 0,0002 | *** |
| Pooling:Sampling_zone | 2 | 2,1705 | 1,0853 | 8,0044 | 0,0008 | *** |
| Residuals | 61 | 8,2706 | 0,1356 | NA | NA |  |

emm contrast results

| Contrast | Pooling | Location | Estimate | SE | df | t.ratio | p.value |
| --- | --- | --- | --- | --- | --- | --- | --- |
| Dried - Frozen | Individual | RhizoCore | 0,0895 | 0,1739 | 61 | 0,5143 | 0,6089 |
| Dried - Frozen | Pooled | Rhisosphere | -0.0529 | 0,1739 | 61 | -0,3044 | 0,7618 |
| Dried - Frozen | Individual | RhizoCore | 0,1480 | 0,1739 | 61 | 0,8512 | 0,3980 |
| Dried - Frozen | Pooled | Rhisosphere | 0,0056 | 0,1739 | 61 | 0,0325 | 0,9742 |
| Dried - Frozen | Individual | RhizoCore | -0,6880 | 0,1791 | 61 | -3,8417 | 0,0003 |
| Dried - Frozen | Pooled | Rhisosphere | -0,8304 | 0,1750 | 61 | -4,7459 | 0,0000 |

### Table S6. Summarized statistics bacteria (A) and fungi (B) beta diversity

| **(A)Bacteria 16S** | | | | | | |
| --- | --- | --- | --- | --- | --- | --- |
| **PERMANOVA** | | | | | | |
| Term | Df | SumOfSqs | R2 | F_value | p_value | Significance |
| Sampling Zone | 2 | 3,50E+13 | 0,317 | 16,694 | 0,001 | ** |
| Preservation | 1 | 2,93E+12 | 0,027 | 2,802 | 0,011 | * |
| Pooling | 1 | 2,08E+14 | 0,019 | 1,986 | 0,045 | * |
| Preservation:Pooling | 1 | 7,16E+14 | 0,06 | 0,684 | 0,762 | ns |
| Zone:Preservation | 2 | 2,68E+14 | 0,024 | 1,281 | 0,184 | ns |
| Zone:Pooling | 2 | 1,99E+14 | 0,018 | 0,95 | 0,484 | ns |
| Residual | 62 | 6,49E+14 | 0,589 | NA | NA | ns |
| Total | 71 | 1,10E+14 | 1 | NA | NA | ns |
| **Permutation test for homogeneity – Sampling Zone** | | | | | | |
| Df | SS | MS | F_statistic | N_Perms | P_value | Significance |
| 2 | 2,06E+14 | 1,03E+14 | 1,33 | 999 | 0,26 | ns |
| 69 | 5,33E+14 | 7,73E+14 | NA | NA | NA |  |
| **Permutation test for homogeneity - Preservation** | | | | | | |
| Df | SS | MS | F_statistic | N_Perms | P_value | Significance |
| 1 | 2,09E+14 | 2,09E+14 | 0,02 | 999 | 0,878 | ns |
| 70 | 7,39E+14 | 1,06E+14 | NA | NA | NA |  |
| **Permutation test for homogeneity - Pooling** | | | | | | |
| Df | SS | MS | F_statistic | N_Perms | P_value | Significance |
| 1 | 1,38E+14 | 1,38E+14 | 15,97 | 999 | 0,002 | ** |
| 70 | 6,05E+14 | 8,64E+14 | NA | NA | NA |  |
| **(B) Fungi ITS** | | | | | | |
| **PERMANOVA** | | | | | | |
| Term | Df | SumOfSqs | R2 | F_value | p_value | Significance |
| Sampling Zone | 2 | 5,41E+14 | 0,367 | 20,786 | 0,001 | ** |
| Preservation | 1 | 1,90E+14 | 0,013 | 1,459 | 0,15 | ns |
| Pooling | 1 | 3,61E+14 | 0,025 | 2,78 | 0,013 | * |
| Preservation:Pooling | 1 | 1,23E+14 | 0,08 | 0,945 | 0,384 | ns |
| Zone:Preservation | 2 | 2,64E+14 | 0,018 | 1,015 | 0,35 | ns |
| Zone:Pooling | 2 | 4,68E+14 | 0,032 | 1,799 | 0,035 | * |
| Residual | 61 | 7,93E+14 | 0,538 | NA | NA | ns |
| Total | 70 | 1,47E+14 | 1 | NA | NA | ns |
| **Permutation test for homogeneity – Sampling Zone** | | | | | | |
| Df | SS | MS | F_statistic | N_Perms | P_value | Significance |
| 2 | 5,25E+13 | 2,63E+14 | 4,39 | 999 | 0,022 | * |
| 68 | 4,07E+14 | 5,98E+14 | NA | NA | NA |  |
| **Permutation test for homogeneity - Preservation** | | | | | | |
| Df | SS | MS | F_statistic | N_Perms | P_value | Significance |
| 1 | 3,25E+08 | 3,25E+08 | 0,00 | 999 | 0,986 | ns |
| 69 | 1,01E+14 | 1,46E+14 | NA | NA | NA |  |
| **Permutation test for homogeneity - Pooling** | | | | | | |
| Df | SS | MS | F_statistic | N_Perms | P_value | Significance |
| 1 | 5,08E+14 | 5,08E+14 | 3,64 | 999 | 0,068 | ns |
| 69 | 9,63E+14 | 1,40E+14 | NA | NA | NA |  |

### Table S7. Statistical assessment of within-group variability in soil microbial community composition across sampling zones, pooling strategies, and preservation methods

| **(A)Bacteria 16S** |  |  |  |  |
| --- | --- | --- | --- | --- |
| Comparison | F_value | P_value | Df | Significance |
| Sampling Zone | 1,33111 | 0,27088 | 2 | ns |
| Pooling Status | 15,96735 | 0,00016 | 1 | *** |
| Preservation Method | 0,01983 | 0,88843 | 1 | ns |
| Pooling within Rhizosphere | 15,01599 | 0,00082 | 1 | *** |
| Pooling within Uncultivated | 75,93097 | 0,00000 | 1 | *** |
| Pooling within RhizoCore | 10,98247 | 0,00316 | 1 | * |

| **(B)Fungi ITS** |  |  |  |  |
| --- | --- | --- | --- | --- |
| Comparison | F_value | P_value | Df | Significance |
| Sampling Zone | 4,39029 | 0,01610 | 2 | * |
| Pooling Status | 3,64034 | 0,06056 | 1 | ns |
| Preservation Method | 0,00022 | 0,98814 | 1 | ns |
| Pooling within Rhizosphere | 43,08438 | 0,00000 | 1 | *** |
| Pooling within Uncultivated | 207,62821 | 0,00000 | 1 | *** |
| Pooling within RhizoCore | 32,17761 | 0,00001 | 1 | *** |

### Table S8. Number of differentially abundant features in bacteria (A) and fungi (B) based of ANCOMBC2 analysis on different taxonomy levels. Summary includes only significant features (adj. p-value> 0,05) with large biological impact (lfc > 1). The color scale from red to green is applied to highlight the changes.

**(A)Bacteria 16S**

|  |  | Differential features on taxonomy level | | | | | |  |
| --- | --- | --- | --- | --- | --- | --- | --- | --- |
| Category | **Comparison** | Phylum | Class | Order | Family | Genus | Species | ASV |
| Pooling | Pooled vs Individual | 0 | 0 | 0 | 1 | 1 | 2 | 11 |
| Preservation | Frozen vs Dried | 1 | 2 | 2 | 3 | 3 | 5 | 27 |
| Sampling zone | RhizoCore vs Rhizosphere | 0 | 0 | 0 | 0 | 1 | 1 | 3 |
| Sampling zone | RhizoCore vs Uncultivated | 8 | 20 | 45 | 72 | 115 | 129 | 482 |
| Sampling zone | Rhizosphere vs Uncultivated | 11 | 31 | 54 | 80 | 131 | 143 | 434 |
| Sampling zone + Pooling | RhizoCore Pooled vs RhizoCore Individual | 0 | 0 | 0 | 0 | 0 | 0 | 0 |
| Sampling zone + Pooling | Rhizosphere Pooled vs Rhizosphere Individual | 0 | 0 | 0 | 0 | 0 | 0 | 0 |
| Sampling zone + Pooling | Uncultivated Pooled vs Uncultivated Individual | 0 | 0 | 0 | 1 | 3 | 4 | 1 |
| Sampling zone + Preservation | RhizoCore Frozen vs RhizoCore Dried | 0 | 3 | 3 | 4 | 3 | 3 | 3 |
| Sampling zone + Preservation | Rhizosphere Frozen vs Rhizosphere Dried | 0 | 1 | 2 | 3 | 3 | 3 | 2 |
| Sampling zone + Preservation | Uncultivated Frozen vs Uncultivated Dried | 0 | 0 | 0 | 0 | 0 | 0 | 0 |

**(B)Fungi ITS**

|  |  | Differential features on taxonomy level | | | | | |  |
| --- | --- | --- | --- | --- | --- | --- | --- | --- |
| Category | **Comparison** | Phylum | Class | Order | Family | Genus | Species | ASV |
| Pooling | Pooled vs Individual | 0 | 0 | 2 | 8 | 13 | 21 | 37 |
| Preservation | Frozen vs Dried | 0 | 1 | 2 | 8 | 18 | 22 | 47 |
| Sampling zone | RhizoCore vs Rhizosphere | 0 | 0 | 1 | 1 | 5 | 7 | 7 |
| Sampling zone | RhizoCore vs Uncultivated | 0 | 10 | 26 | 46 | 68 | 79 | 73 |
| Sampling zone | Rhizosphere vs Uncultivated | 0 | 10 | 24 | 45 | 72 | 82 | 77 |
| Sampling zone + Pooling | RhizoCore Pooled vs RhizoCore Individual | 0 | 0 | 1 | 1 | 2 | 1 | 0 |
| Sampling zone + Pooling | Rhizosphere Pooled vs Rhizosphere Individual | 0 | 1 | 0 | 0 | 0 | 0 | 0 |
| Sampling zone + Pooling | Uncultivated Pooled vs Uncultivated Individual | 0 | 0 | 2 | 3 | 4 | 2 | 2 |
| Sampling zone + Preservation | RhizoCore Frozen vs RhizoCore Dried | 0 | 0 | 0 | 0 | 0 | 0 | 0 |
| Sampling zone + Preservation | Rhizosphere Frozen vs Rhizosphere Dried | 0 | 1 | 2 | 1 | 1 | 1 | 3 |
| Sampling zone + Preservation | Uncultivated Frozen vs Uncultivated Dried | 0 | 0 | 0 | 1 | 1 | 2 | 0 |
